# The bodily appearance of a virtual partner affects the activity of the action observation and action monitoring systems in a minimally interactive task

**DOI:** 10.1101/2024.07.03.601841

**Authors:** U.G. Pesci, Q. Moreau, V. Era, M. Candidi

## Abstract

One pending question in social neuroscience is whether interpersonal interactions are processed differently by the brain depending on the bodily characteristics of the interactor, i.e., their physical appearance. To address this issue, we engaged participants in a minimally interactive task with an avatar either showing bodily features or not while recording their brain activity using Electroencephalography (EEG) in order to investigate indices of action observation and action monitoring processing. Multivariate results showed that bodily compared to non-bodily appearance modulated parieto-occipital neural patterns throughout the entire duration of the observed movement and that, importantly, such patterns differ from the ones related to initial shape processing. Furthermore, among the electrocortical indices of action monitoring, only the early observational Positivity (oPe) was responsive to the bodily appearance of the observed agent under the specific task requirement to predict the partner movement. Taken together, these findings broaden the understanding of how bodily appearance shapes the spatiotemporal processing of an interactor’s movements. This holds particular relevance in our modern society, where human-virtual agent interactions are rapidly becoming ubiquitous.

## INTRODUCTION

Observing and monitoring others’ actions are essential functions for humans’ social life and support interpersonal coordination (Cardellicchio et al., 2021; Knoblich & Sebanz, 2008; Ridderinkhof et al., 2004; Vesper et al., 2010). Neuroimaging studies have identified, among others, two brain networks involved in the prediction and monitoring of observed actions, namely the Action Observation Network (AON, Grafton & de C. Hamilton, 2007), and the action monitoring system (AMS, Cohen, 2016; De Bruijn et al., 2007; Malfait et al., 2010; Miltner et al., 2004; Ridderinkhof et al., 2004; van Schie et al., 2004).

The AON comprises parietal (anterior IntraParietal Sulcus, IPS), premotor (ventral and dorsal PreMotor, vPM, dPM), motor (M1) and visual (lateral occipito-temporal cortex, LOTC) areas (Hardwick et al., 2018). Within the LOTC, the Extrastriate Body Area (EBA) is thought to be specifically involved in human body form processing (Downing et al., 2001; Urgesi et al., 2006; Candidi et al., 2007; Pitcher et al., 2009, Fusco et al., 2022) and the Superior Temporal Sulcus (STS) is thought to be crucial for the processing of dynamical aspects of visual motion, (i.e., biological motion, (Allison et al., 2000; Lingnau & Downing, 2015; Pitcher & Ungerleider, 2021; Puce & Perrett, 2003; Saygin et al., 2012). EEG studies have identified the alpha/mu desynchronization (8-13 Hz power range) over centro-parietal electrodes as the main EEG marker of the AON activity, since it occurs during action observation and is similar to the markers of action execution (Cochin et al., 1999; Pineda, 2005). Alpha/mu desynchronization has also been considered to reflect the activity of the so-called Mirror Neuron System (MNS - (Rizzolatti & Craighero, 2004). Nevertheless, recent results challenge a dependency of the alpha/mu suppression only by the MNS activity, suggesting that the specificity of this activity to somatosensory features of observed and executed actions might be related to sensory processing rather than motor mirroring (Coll et al., 2017).

The AMS has been described extensively for its role in the control of executed actions, and has been also associated to the detection of sudden and/or unexpected changes in the individual actions or in those of others (e.g., a change in the trajectory of an observed movement; (De Bruijn et al., 2007; Malfait et al., 2010; Miltner et al., 2004; Moreau et al., 2020, 2022; Shane et al., 2008; Spinelli et al., 2018; van Schie et al., 2004). This system is involved in a constant process of prediction and adaptation (Ullsperger, 2017; Ullsperger et al., 2014; Wilken et al., 2023) and comprises the posterior part of the medial frontal cortex (pMFC), specifically the anterior cingulate cortex (ACC), the pre-supplementary motor area (pre-SMA) and the adjacent dorsomedial prefrontal cortex (dmPF, (Nee et al., 2011; Ridderinkhof et al., 2004). Although these areas often show complex and overlapping patterns of activation, the ACC is considered to be the central hub of the action monitoring system and one of the main sources of the electrocortical indices of action monitoring. Specifically, previous EEG studies identified two event-related potentials (ERPs) linked to action monitoring processes, the error-related negativity (ERN; (Falkenstein et al., 1991; Gehring et al., 1993; Pezzetta et al., 2018, 2023; Spinelli et al., 2018; Taylor et al., 2007) and the error positivity (Pe; (Falkenstein et al., 2000). These ERPs are associated with the detection of both errors committed by the self and errors observed in others. In the latter case they are referred to as observation-ERN (oERN) and observation-Pe (oPe; Spinelli et al., 2018; van Schie et al., 2004). Furthermore, action monitoring is indexed in the time-frequency domain by the so-called midfrontal Theta rhythm (Cohen, 2011), a synchronization in the Theta band (4-8 Hz) over midfrontal sites (i.e. electrode FCz) that tends to last a few hundred milliseconds (200 to 400) after an error is detected and can be easily observed in individual subjects. Both time and time-frequency domain indexes have been observed during interpersonal motor interactions (Moreau et al., 2020, 2022). In detail, recent studies have shown that the amplitude of the ERN and the Pe, as well as of midfrontal Theta’s power synchronization are modulated depending on the social context where prediction errors occur, with a generally stronger response during highly interactive tasks (i.e. tasks requiring a continuous monitoring of others’ actions) compared to less-to-not interactive tasks (i.e. tasks where the subject’s behaviour was independent from others’ actions; (Moreau et al., 2020, 2022).

One pending question concerning the interaction between action observation/monitoring and interpersonal motor coordination is whether biological visual motion can induce activity in the AON and AMS independently from the visual appearance of the observed/monitored agent. This is crucial to better understand the neural system supporting social observation as well as dynamic social interaction processes where reciprocal and mutual adaptation are involved (Tidoni et al., 2022). Previous studies have manipulated the physical appearance of observed bodies and /or the kinematics of the movements (Albertini et al., 2021; Craighero et al., 2008; Gazzola et al., 2007; Hofree et al., 2015; Kupferberg et al., 2018; Miyamoto et al., 2023; Perani et al., 2001; Shimada, 2010; Tai et al., 2004) but results remain elusive and inconsistent across paradigms and experimental techniques (i.e., EEG, fMRI). Furthermore, to the best of our knowledge, none of them investigated the modulation of those processes, both at the behavioural and neural level, in dynamic social contexts.

Hence, overall, to what extent the AON is triggered by motion perception without the concurrent appearance of a full body form is still a matter of debate. Since recent results have also shown that Theta oscillatory activity in occipito-temporal nodes of the AON (i.e., LOTC) are in phase with the activation of the action monitoring system during social (human-avatar) interactions (Moreau et al., 2020) we also aim at investigating if and how the appearance of an interactor during online (minimal) interactions affect the neural patterns related to action monitoring. In order to do so, we engaged human participants in a minimally interactive version of a visuo-motor interpersonal interaction task (Candidi et al., 2017; Era et al., 2023; Sacheli et al., 2015). Our first aim focused on the AON to investigate 1) whether the alpha/mu desynchronization over centro-parietal electrodes was modulated by the appearance of the interactor (Cochin et al., 1999; Pineda, 2005; Tucciarelli et al., 2015) compared to occipital alpha/mu activity, and 2) explore time-locked data with multivariate pattern analysis (MVPA) to examine if and at which point in time the brain distinguishes bodily from non-bodily movements. Our second aim focused on replicating and broadening previous results on the activity in the action monitoring system hypothesizing a higher amplitude of the oERN and of the oPe and a stronger synchronization of midfrontal Theta in response to corrections in the trajectory performed by an avatar compared to a set of dots.

## METHODS

### Participants

21 participants took part in the experiment. The sample size was based on a power analysis, performed with the software More Power (Campbell & Thompson, 2012). More specifically, we indicated as expected effect size the partial eta squared value obtained by Moreau et al. (2020) (i.e., .33). The output indicates that a 2 x 2 x 2 within subject design, a power of .80 and a partial eta squared of .33, requires a sample size of 18 participants. All participants were right-handed with normal or corrected-to-normal vision. Participants were naive as to the aim of the experiment and were informed of the purpose of the study only after all the experimental procedures were completed. One participant was removed from all results due to technical issues during the recording. Thus, the final sample includes 20 participants (10 females, mean age: 24; S.D. = 2.5). Participants signed informed consents approved along with all experimental procedures by the Ethics Committee of the Fondazione Santa Lucia (Rome, Italy), approval number Prot. CE/prog 835, and the study was performed in accordance with the 2013 Declaration of Helsinki..

### Experimental stimuli and set-up

Participants sat in front of a desk and observed visual stimuli on a 1.024 x 768 resolution LCD monitor placed at ∼60 cm from their eyes. Using a keyboard connected to the PC, participants were asked to press a button (up/down arrow) with the index/middle finger of their right hand at the exact same time when an virtual partner (VP - facing the participant) on the screen, or two dots in a control condition, touched a bottle-shaped object in front of him. The bottle-shaped object is constituted of two superimposed cylinders of different diameters (see below for details on task). The variability of the appearance of the interactor allowed us to explore the influence on the EEG correlates of the presence of a body on action observation.

The keyboard’s press reaction times were recorded using E-Prime 2 Professional software (Psychology Software Tools Inc., Pittsburgh, PA). The VP’s index-thumb grasping time was measured trial-by-trial by means of a photodiode placed on the screen sending a signal recorded by means of a TriggerStation (BrainTrends ltd., Italy). The photodiode was triggered by a white dot displayed on the screen (not visible to the participants) during the clip frame corresponding to the instant when the avatar grasped (or the dots reached) the virtual object.

Video stimuli were adapted from Gandolfo *et al*. (Gandolfo et al., 2019) in which reaching and grasping movements of a virtual character are based on the trajectories of real human actors performing a number of grasping movements (Figure 1). As a control condition, the same motion kinematics of the human shoulders, right index and thumb were implemented in a set of four dots connected by lines in a manner aimed at reducing the perception of the shape of a human body and create a set of non-humanoid stimuli (hereafter NonBody stimuli) (Figure 1. Kinematic features of the actors were recorded using a SMART-D motion capture system (MoCAP) [Bioengineering Technology & Systems (B|T|S)] attaching infrared reflective markers (5 mm diameter) to an actor reaching and grasping with his right hand a cylinder on a table. Four infrared cameras with wide-angle lenses placed about 100 cm away from each of the four corners of the table captured at 100 Hz the movement of the markers in the 3D space (Tieri et al., 2015). The kinematics were eventually transferred on the virtual VP’s bones by means of Motion Builder 2011 (Autodesk, Inc.).

**Figure 1.**
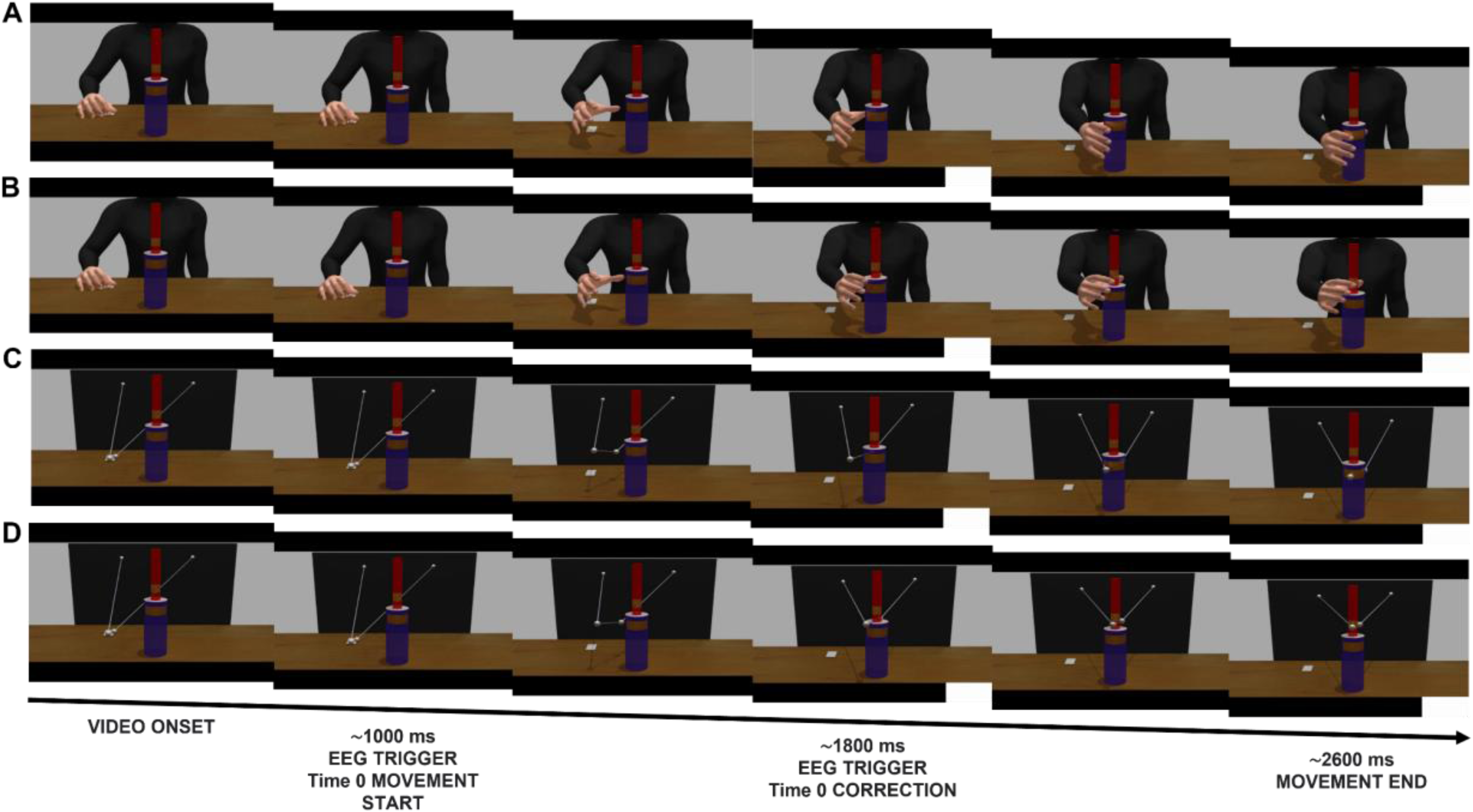
Examples of the sequence of frames for each combination of the Appearance Factor with the Correction factor: A) Body - NoCorrection; B) Body - Correction; C) NonBody - NoCorrection; D) NonBody - Correction.

Stimuli were thus divided equally in two main categories representing: 1) a humanoid partner (“Body”, Figure 1a/b), applying the trajectories to a Caucasian male character, and 2) a non-humanoid stimulus (“NonBody”, Figure 1c/d), composed of four spheres representing the right and left shoulder, thumb and index finger of the right hand, connected by white lines. Note that the character was presented only with the upper body from the shoulders down, since variables related to face were to be ignored by participants. The complete sample of clips comprised twelve different grasping movements (Grip factor), half of them ending with a precision grip (grasping the top part of the bottle-shaped object), and half ending with a power grip (on the bottom part). Each movement had a version with the Body and one with the NonBody (Appearance Factor). The Appearance Factor is fundamental for the present purposes as it represents the main manipulation of the stimuli to test the role of physical appearance of a human body in triggering behavioural and neural resources associated to action observation, prediction, and error-processing.

Crucially for the hypothesis regarding action monitoring, in 30% of the trials the movements included an online correction of trajectory by the VP or the dots (Correction Factor - Fig. 1 B/D), which switched from a precision to a power grip (or vice versa) during the reaching phase at different times for each video. In order to lock on the EEG signal the exact time of this correction, the photodiode was triggered by another white dot, identical to the abovementioned one, flashed during the clip frame corresponding to the start of the online correction, or in the corresponding time in the NoCorrection trials.

### Experimental task

Participants were asked to press a button corresponding either to the upper part of the bottle-shaped object (up-arrow) or the bottom part of it (down-arrow) as synchronously as possible with the touch time of the VP or of the dots on the bottle in the virtual environment. Participants had to follow an auditory instruction concerning the response to be executed, which was delivered prior to each trial via headphones. Specifically, interactions occurred in two different conditions (Interactivity Factor): 1) a Cued condition, where participants knew in advance which key they had to press and therefore only had to track the timing of the VP’s/dots’ grasping time; 2) an Interactive condition, where they did not know which key they had to press and had to adapt to the movement of the VP/dots, therefore having to anticipate and monitor it online to coordinate not only temporally but also spatially with it.

In detail, instructions for the Cued condition consisted of either a high pitch (indicating that participants had to press the upper arrow on their keyboard) or a low pitch sound (indicating that participants had to press the lower arrow on their keyboard). For the Interactive condition, instructions consisted in a sound saying “Ugua”, shortened for Italian “Uguale” (“Same”), asking the subjects to press the key matching the VP’s behaviour or dots contact site (e.g. if the avatar grasped, or the dots reached, the upper part of the object, participant had to press the up arrow) or “Oppo”, shortened for Italian “Opposto” (“Opposite”), asking the subjects to press the key opposite to the VP’s behaviour or dots contact site (e.g. if the avatar grasped, or the dots reached, the upper part of the object, participant had to press the down arrow). At the end of each trial, participants received feedback on their performance according to both synchrony (ms) and correct pressing site (according to the instruction received at the beginning).

The task presented above is a revision of the ecological and controlled human-avatar “Joint Grasping” task (Candidi et al., 2017; Era et al., 2020; Moreau et al., 2020; Sacheli et al., 2018), which has been shown to recruit similar behavioural and neural processes called into play during human-human interaction (Candidi et al., 2015; Era et al., 2018).

Participants performed in total eight 100-trial blocks (4 blocks of the Body condition, 4 for the Non-Body condition, each divided in two successive Interactive and two successive Cued ones, presented in a counterbalanced order between participants). Stimuli presentation and randomization were controlled by E-Prime 2 Professional Software (Psychology Software Tools Inc.).

### Behavioural Data

The Interpersonal Synchrony with the VP’s/dots’ grasping was considered as the main behavioral measure, computed as the absolute value of the time delay between the participant’s pressing of the keyboard button and the VP’s/dots’ bottle touch time. This is believed to show the success of human-avatar interaction achieved by monitoring and predicting the movements of the stimulus. At the group level, we checked for the presence of outliers in our sample by investigating whether any of our participants’ Interpersonal Synchrony across all conditions was 2.5 standard deviations above the group mean. According to this control, all 20 subjects were included in the final analyses.

### EEG recording and preprocessing

EEG signals were recorded and amplified using a Neuroscan SynAmps RT amplifiers system (Compumedics Limited, Melbourne, Australia). These signals were acquired from 58 tin scalp electrodes embedded in a fabric cap (Electro-Cap International, Eaton, OH), arranged according to the 10-20 system. The EEG was recorded from the following channels: Fp1, Fpz, Fp2, AF3, AF4, F7, F5, F3, F1, Fz, F2, F4, F6, F8, FC5, FC3, FC1, FCz, FC2, FC4, FC6, T7, C5, C3, C1, Cz, C2, C4, C6, T8, TP7, CP5, CP3, CP1, CPz, CP2, CP4, CP6, TP8, P7, P5, P3, P1, Pz, P2, P4, P6, P8, PO7, PO3, AF7, POz, AF8, PO4, PO8, O1, Oz and O2. Horizontal electro-oculogram (HEOG) was recorded bipolarly from electrodes placed on the outer catchi of each eye and signals from the left earlobe were also recorded. All electrodes were physically referenced to an electrode placed on the right earlobe and were algebraically re-referenced off-line to the average of both earlobe electrodes. Impedance was kept below 5 KΩ for all electrodes for the whole duration of the experiment, amplifier hardware band-pass filter was 0.1-200 Hz and sampling rate was 1000 Hz. To remove ocular artifacts (eye blinks and saccades), a blind source separation method, Independent Component Analysis (ICA) (Jung et al., 2000), was applied on continuous raw signal. Artefactual components were removed based on the topography and the explained variance, and data were visually inspected to check the efficient removal of blinks and saccadic movement correlates from the EEG signal. The signal was then low-pass filtered at 45 Hz, and segmented into epochs of 8000 ms around the trigger corresponding to the start of the video (Go - trigger). To investigate the neural patterns related to action observation or action monitoring, the time 0 was manually moved to either the frame when the movement of the VP or of the dots started (monitor t0) or to the frame when a correction occurred (or the equivalent one in No-Correction trials - error t0), respectively. Segmented data were visually inspected and remaining trials containing ocular and muscular artifacts as well as trials with 0 behavioural accuracy were removed from further analysis. The average number of trials per participant after artefact rejection was 351 trials for Body (with 39 Interactive-Correction, 127 Interactive NoCorrection, 59 Cued-Correction, 126 Cued-NoCorrection) and 347 trials for NonBody (with 39 Interactive-Correction, 121 Interactive NoCorrection, 59 Cued-Correction, 128 Cued-NoCorrection). Importantly, in the analysis of action monitoring activity, the epochs were restricted to −2000 ms to +2000 ms around the trigger corresponding to the VP/dots’ Correction/NoCorrection frame. All preprocessing and further univariate analyses were performed using the FieldTrip (version 2019-04-03) routines (Donders Institute, Nijmegen; (Oostenveld et al., 2011) in Matlab R2019a (The MathWorks, Inc.).

### EEG Analysis

#### Univariate Analysis

In order to investigate whether the neural correlates of action monitoring would be modulated by the appearance of the interacting partner (Appearance factor) we analysed the event-related potentials (ERPs) over fronto-central electrodes (i.e., FCz) time-locked to the correction of the observed movement’s trajectory (error t0), as well as the activity in the time-frequency domain focused on midfrontal Theta. Moreover, in order to investigate whether the appearance of the interacting partner would modulate earlier neural processes related to action observation, we also analysed the alpha/mu desynchronization over centro-parietal electrodes (Cochin et al., 1999; Pineda, 2005; Tucciarelli et al., 2015) time-locked to the start of the observed movement (movement t0).

##### ERPs

Before ERP averaging across subjects for each condition, EEG time-series were high-pass filtered at 1 Hz to remove slow drifts from the data, hence reducing the contribution of slow potentials. Since we found no oERN in response to the observation of a partner’s correction in any condition (see Discussion), only the early oPe component was further analysed being quantified as the mean amplitude in the time window between 250-400 ms over electrode FCz.

##### ERD/S

After preprocessing, as in the time-lock analysis, we high-pass filtered the segmented EEG signal at 1 Hz. Moreover, to remove the effects of ERPs in the time-frequency domain we subtracted the mean evoked response from each single trial, thus removing phase-locked activity (Sauseng et al., 2007). Each epoch was then put in the frequency domain using Hanning-tapered window (Cohen, 2014) with a 50 ms time resolution, obtaining single-trial power representations for frequencies in a range from 1 to 40 Hz in steps of 1. The obtained induced power was averaged over trials for each condition and each subject, and the grand averages across subjects were displayed as event-related desynchronization/synchronization (ERD/ERS) with respect to a baseline period ranging from -.2 to 0 ms before the correction occurred (for the error-locked analysis), and from −1300 to −1000 ms before the start of the movement (for the action observation analysis). Only the epochs when the virtual partner did not correct its trajectory (NoCorrection trials) were included in the action observation analysis.

For midfrontal Theta, we extracted the ERD/ERS for the Theta band (3-7 Hz) between 200 and 500 ms after the correction (or the corresponding frame in the NoCorrection trials) and analysed the modulation of power over FCz. For the action observation analysis, we extracted the ERD/ERS for the alpha (8-13 Hz) band from 0 to 1600 ms (average movement time) over centro-parietal electrodes (CPz/1/2/3/4). As a control analysis (Coll et al., 2017), we also extracted the ERD/ERS in the same frequency band and time window over an equal number of occipital electrodes (Oz/1/2/PO7/8).

#### Multivariate analysis

To better identify the neural patterns of action observation that were sensitive to the appearance of the observed interactor (i.e., a Body or a NonBody), we adopted a multivariate approach. Multivariate analyses are more sensitive than univariate analyses (Haxby et al., 2001; Tucciarelli et al., 2015), since they assume that the processing of different stimulus category has different neural patterns associated to be exploited, thus they treat whole-brain sensor/source-level data as response patterns rather than investigating changes between average responses for each single sensor/source (as in the univariate approach) (Grootswagers et al., 2017a). Analyses were performed using the MVPA-Light toolbox (Treder, 2020) in Matlab R2019a. Prior to every classification, data were resampled at 250 Hz and normalized using z-scores to center and scale the training data, providing numerical stability (Treder, 2020). Across all classification analyses, if a class had fewer trials than another, we corrected the imbalance by undersampling the over-represented condition (i.e., randomly removing trials). Then, using the preprocessing option mv_preprocess_average_samples, training data (i.e., EEG trials) from the same class (i.e., Body/NonBody) were randomly split into 5 groups and averaged, so that the single-trial dataset was replaced by 5 averaged trials (i.e., samples). The classifications were then run using these averaged samples, as this step is known to increase the signal-to-noise ratio (Grootswagers et al., 2017b; Smith & Smith, 2019; Treder, 2020).

##### Classification across time

We first performed a binary classification analysis in time (- 1200 to 2000 ms around movement start), using Body/NonBody samples. To do so, a Linear Discriminant Analysis (LDA) classifier was implemented using the MVPA-Light toolbox. For linearly separable data, an LDA classifier divides the data space into *n* regions, depending on the number of classes, and finds the optimal separating boundary between them using a discriminant function to find whether the data fall on the decision boundary (i.e., 50% classification accuracy) or far from it (i.e., > 50% classification accuracy). A k-fold cross-validation procedure in which samples (5 x 2 classes = 10 total samples) were divided into 10 folds was repeated 10 times so that each sample was either used for training or testing at least once, and the classifier’s % accuracy (acc) was used to assess decoding performance over time in each comparison. Statistical significance (i.e., performance significantly above 0.5 chance level) was assessed through non-parametric cluster-based permutation tests (Maris & Oostenveld, 2007), using ‘maxsum’ for cluster corrections and Wilcoxon test for significance assessment.

In order to further understand how the different patterns related to observing Body and NonBody motion evolve over time, we adopted the temporal generalization method (King & Dehaene, 2014). Specifically, we tested whether the neural representations carrying the classifier performance were stable (generalizable) across time, or rather they differed. In order to do so, we trained our classifier on each single time point and tested it at all time points in the same time window previously selected for the classification across time. This provided a temporal generalization matrix (time x time) with accuracy values tested for statistical significance.

Moreover, to investigate what electrodes contributed most to the classification over time, we performed a searchlight analysis using two neighboring matrices for time points and electrodes, respectively. Searchlight analysis is one approach to localize multivariate effects, as it strikes a balance between localization and statistical power (Kriegeskorte et al., 2006; Treder, 2020). Thus, in this analysis each electrode/time-point and its direct neighbors acted as features for the classification, resulting in a channels x time points matrix of accuracy scores tested for statistical significance. By plotting the results from this matrix also on a spatial topography averaged in specific time windows of interest, we then visualized which electrodes carried the most weight in the temporal decoding.

Lastly, we were interested in finding whether the neural patterns related to Body and NonBody motion were classifiable under different interactive scenarios (i.e., in the Interactive condition compared to the Cued one). Thus, we performed a multiclass Linear Discriminant Analysis (LDA) in time to classify the EEG activity in the time window from movement onset (0 s) to its average end (1600 ms) for each of the four classes (Body - Interactive, Body - Cued, NonBody - Interactive, NonBody - Cued). The accuracy values across testing folds of all repetitions were then averaged and presented on a confusion matrix to assess the probability of the classifier to in/correctly assign classes.

### Data handling and statistics

Interpersonal Synchrony values, error-locked ERP mean amplitudes in the oPe time window (250-400 ms) and midfrontal Theta synchronization values were analysed through a 2×2×2, within subject, repeated measures ANOVA with Appearance (Body/NonBody), Interactivity (Interactive/Cued), and Correction (Correction/NoCorrection) as within-subject factors. Conversely, while our primary interest concerning movement-locked time-frequency induced ERD/ERS focused on the interaction between the factors Interactivity and Appearance, we initially analysed it by also including a factor Cluster (Occipital/Centro-Parietal) through a 2×2×2, within subject, repeated measures ANOVA with Appearance (Body/NonBody), Interactivity (Interactive/Cued) and Cluster (Occipital/Centro-Parietal) as within-subject factors. After this first exploratory analysis aimed at testing the topographical distribution of the effects of Appearance and Interactivity, movement-locked time-frequency induced ERD/ERS values were analysed through two separate 2x2, within subject, repeated measures ANOVAs with Appearance (Body/NonBody) and Interactivity (Interactive/Cued) as within-subject factors focusing, respectively, on the activity over five centro-parietal electrodes and on the activity over five control occipital electrodes (similarly to (Coll et al., 2017). Since our main hypothesis did not focus on Interaction Type (Complementary/Imitative) and Movement Type (Up/Down), these factors were collapsed in order to have a higher number of trials for each condition. Normality and homoscedasticity assumptions were checked using the Shapiro-Wilk test, revealing that, only for behavioural data, NonBody data were not normally distributed compared to Body ones.

All statistical analyses were performed in Matlab R2019a, JASP Software and R using the DABEST Package (Ho et al., 2019). Post-hoc correction for multiple comparisons ((Ryan, 1959)) was conducted applying the Bonferroni method (Maris & Oostenveld, 2007) to the interactions that resulted as significant (*p.* <.05) from the ANOVA.

## RESULTS

### Behavioural

The 2 Appearance (Body/NonBody) x 2 Interactivity (Interactive/Cued) x 2 Correction (Correction/NoCorrection) ANOVA showed that Interpersonal Synchrony was significantly modulated by the Correction factor (Figure 2A) [F(1, 19) = 39.71, p <.001, ηp² = .676]. Specifically, Interpersonal Synchrony was worse when the stimulus (either virtual-partner/dots-pattern) did not correct its trajectory (M = 127.40 ms, SD = 41.83) rather than when a correction occurred (M = 106.22 ms, SD = 29.05).The analysis also showed a significant interaction between the Appearance and Correction factors [F(1, 19) = 14.32, p = .001, ηp² = .43]. Post-hoc tests indicated that Interpersonal Synchrony was worse in the NonBody-NoCorrection condition compared to the NonBody-Correction one (p < .001) and the Body-NoCorrection one (p = .020). Also the interaction between the Interactivity and the Correction factor resulted significant [F(1, 15) = 13.07, p = .002, ηp² = .41], with interpersonal Synchrony being worse in NoCorrection trials compared to Correction trials both in the Free (p < .001) and in the Cued (p = .020) condition (Figure 2B). These results suggest that, independently from the level of interactivity implied in the condition (i.e., Interactive or Cued), a correction in the trajectory of the observed movement might have prompted subjects’ attention, improving their ability to predict VP’s/dots’ reaching time. Regarding the Appearance factor, we observed that the appearance of the interactor seemed to improve the performance of the subjects only during NoCorrection trials, i.e., when subjects’ performance was generally worse, compared to the Correction ones.

**Figure 2.**
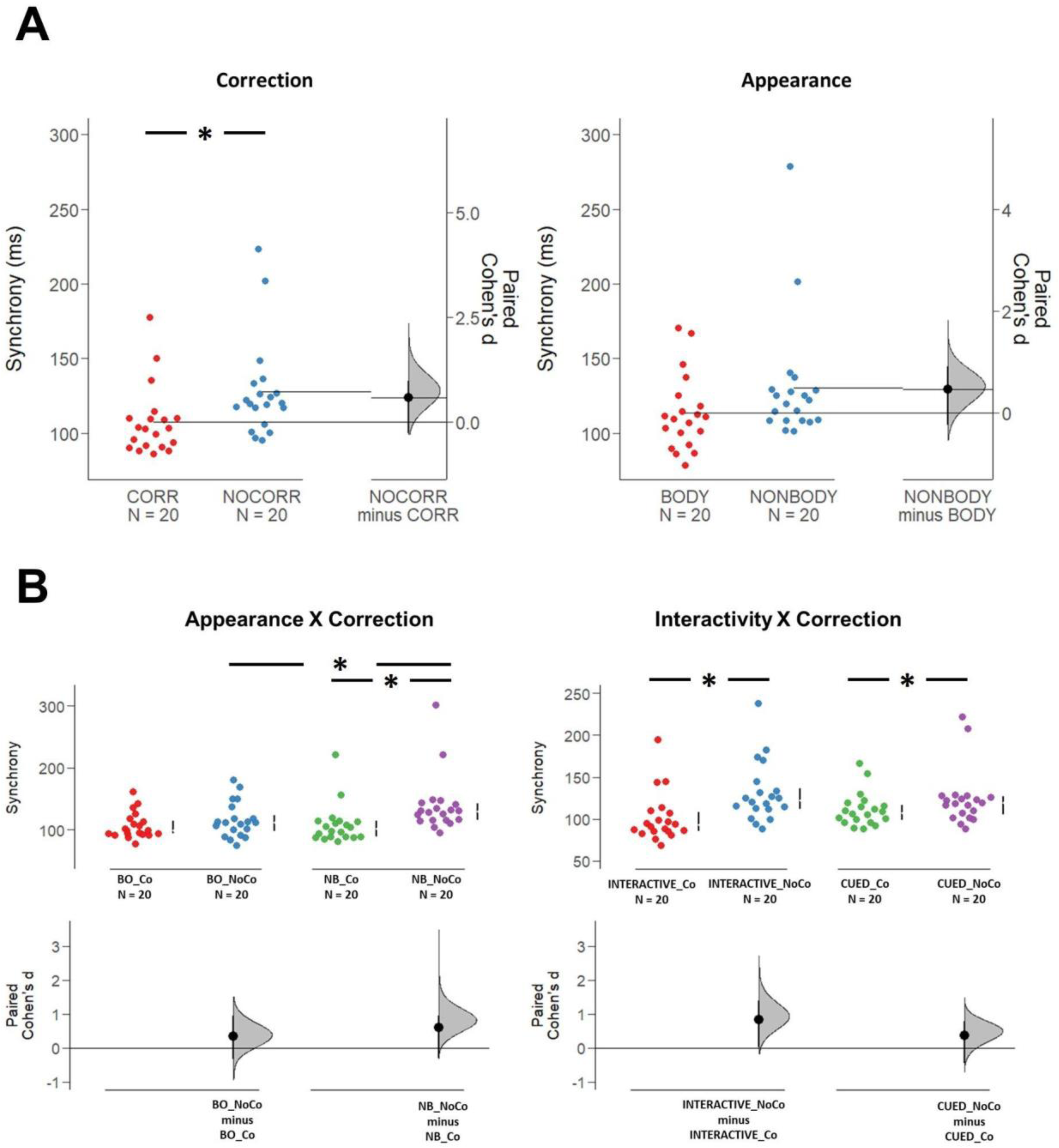
Top row, left panel: significant modulation of Interpersonal Synchrony values by the Correction Factor (left panel - F(1, 19) = 39.71, p <.001, ηp² = .676). Right panel: nearly significant modulation of Interpersonal Synchrony by the Appearance factor (right panel - F(1, 19) = 3.61, p = .07, ηp² = .160). Bottom row, left panel: significant interaction between Appearance and Correction factor (F(1, 19) = 14.32, p = .001, ηp² = .43). The post-hoc test indicated that Interpersonal Synchrony was worse in the NonBody-NoCorrection condition compared to the NonBody-Correction one (p < .001) and the Body-NoCorrection one (p = .020). Right Panel: significant interaction between Interactivity and Correction factor (F(1, 15) = 13.07, p = .002, ηp² = .41). The post-hoc test indicated that Interpersonal Synchrony was worse in NoCorrection trials compared to Correction trials both in the Free (p < .001) and in the Cued (p = .020) condition.

### EEG results

#### ERPs - oPe

The 2 Appearance (Body/NonBody) x 2 Interactivity (Interactive/Cued) x 2 Correction (Correction/NoCorrection) ANOVA on oPe amplitudes revealed a main effect of the Correction factor [F(1, 19) = 32.119, p <.001, ηp² = .628] showing that the early oPe amplitude was larger for Correction (M = 1.015, SD = 0.86) compared to NoCorrection trials (M = 0.127, SD = 0.192). There was also a significant main effect of Interactivity [F(1, 19) = 18.438, p <.001, ηp² = .492], with larger oPe for Interactive (M = 0.745, SD = 0.95) compared to Cued trials (M = 0.398, SD = 0.46). oPe amplitude was also larger for Body (M = 0.626, SD = 0.86) compared to NonBody trials (M = 0.517, SD = 0.66), as the Appearance factor approaches ignificance threshold [F(1, 19) = 4.298, p = .052, ηp² = .184]. The three factors interacted significantly [F(1, 19) = 8.787, p = .008, ηp² = .316]. Post-hoc tests indicated that the early oPe amplitude was significantly larger during Body-Interactive-Correction trials than during Body-Interactive-NoCorrection trials (*p* < .001), Body-Cued-Correction trials (*p* < .001) and NonBody-Interactive-Correction trials (*p* < .001). Moreover, the early oPe amplitude recorded during NonBody-Interactive-Correction trials was larger than during NonBody-Interactive-NoCorrection trials (*p* < .001), as well as than during NonBody-Cued-Correction trials (*p* = .010) (see Figure 3). This pattern of results is in line with previous results (Moreau et al., 2020) showing a modulation of the Pe amplitude by higher-order task-related factors (Interactivity factor), and they add new information regarding the electrophysiological indices of action monitoring when observing a bodily movement compared to a non-bodily one (Appearance factor).

**Figure 3.**
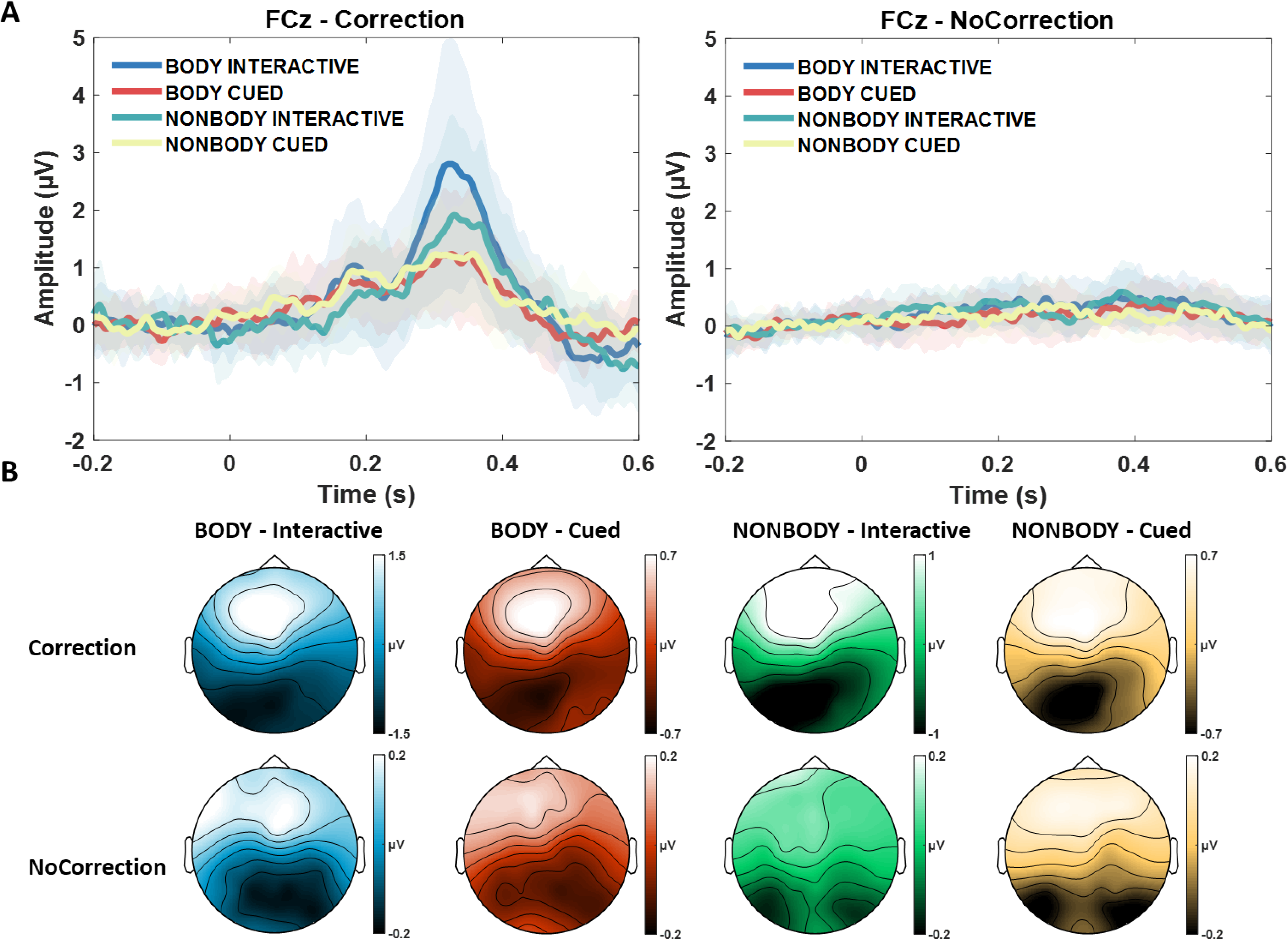
A) Grand averages of the early oPe component over FCz in all experimental conditions with (left panel) and without (right panel) correction. Data are time-locked to the correction of the Body/dots (or the equivalent frame when no correction occurred). B) Topographies of the early oPe component (from 250 to 400 ms after the correction of the observed movement’s trajectory) for each condition.

### Time frequency analyses - midfrontal Theta, and centro-parietal alpha/mu de/synchronization

#### Correction-locked midfrontal Theta over central electrodes

The 2 Appearance (Body/NonBody) x 2 Interactivity (Interactive/Cued) x 2 Correction (Correction/NoCorrection) ANOVA showed a main effect of Correction, with higher Theta synchronization for Correction (M = 0.564, SD = 0.479) compared to NoCorrection trials (M = 0.046, SD = 0.138) [F(1, 19) = 59.822, p <.001, ηp² = .759]. Moreover, the Interactivity factor reached statistical significance, with larger Theta synchronization during Interactive (M = 0.417, SD = 0.528) compared to Cued trials (M = 0.193, SD = 0.283) [F(1, 19) = 19.790, p <.001, ηp² = .510]. The Correction and Interactivity factors interacted significantly [F(1, 19) = 12.365, p <.002, ηp² = .394]. Post-hoc tests indicated that Theta synchronization was larger for Interactive-Correction trials compared to all other condition (all *ps*. < .001) as well as for Cued-Correction trials compared to Cued-NoCorrection trials (*p.* <. 001). These results are in line with previous ones from our group (Moreau et al., 2020), but contrary to our hypothesis, no statistical difference in Theta synchronization was found between Body and NonBody trials (see Figure 4).

**Figure 4.**
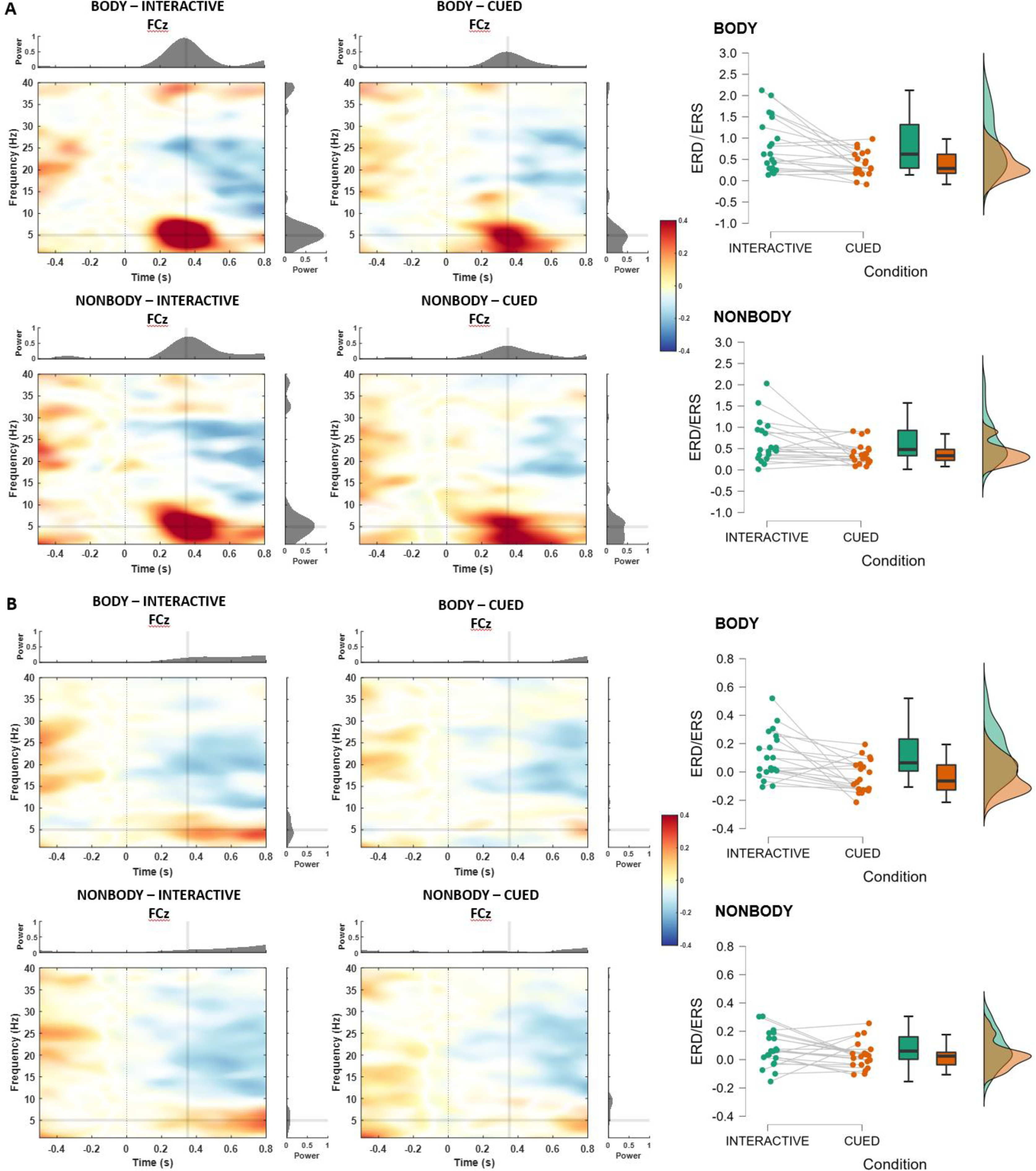
Power change relative to baseline over electrode FCz for all conditions. A) time-frequency representations for all the conditions where a correction occurred. B) Time-frequency representations for all the conditions with no correction (time 0 represents the frame equivalent to the one when the movement changed during the correction trials). Raincloud plots on the right show the mean Theta ERS over FCz in the time window between 200 and 500 ms.

#### Movement-monitoring power over centro-parietal electrodes

To investigate the alpha/mu (8-13 Hz) desynchronization related to action observation over centro-parietal electrodes, compared to visual alpha desynchronization over occipital electrodes as control, we run a 2 Cluster (Occipital/Centro-Parietal) x 2 Appearance (Body/NonBody) x 2 Interactivity(Interactive/Cued), within subjects, repeated measures ANOVA. Both the Cluster [F(1, 19) = 38.324, p < .001, ηp² = .668] and the Interactivity factor [F(1, 19) = 5.574, p = .029, ηp² = .227] reached statistical significance but their interaction did not [F(1, 19) = 2.586, p = .124, ηp² = .120]. Thus, we further conducted two separate 2 Appearance x 2 Interactivity ANOVAs focusing on alpha/mu desynchronization over, respectively, only the centro-parietal cluster or the occipital cluster. As shown in Figure 5A, over centro-parietal electrodes, the Interactivity factor reached statistical significance, with stronger alpha desynchronization during Interactive (M = −0.287, SD = 0.171) compared to Cued trials (M = −0.224, SD = 0.147) [F(1, 19) = 5.233, p = .034, ηp² = .216]. Similar to the midfrontal Theta results, no significant difference was found between the alpha desynchronization during the observation of Body and NonBody movements [F(1, 19) = 0.198, p = .662, ηp² = .010] and no factor reached significance in the same analysis over occipital electrodes.

**Figure 5.**
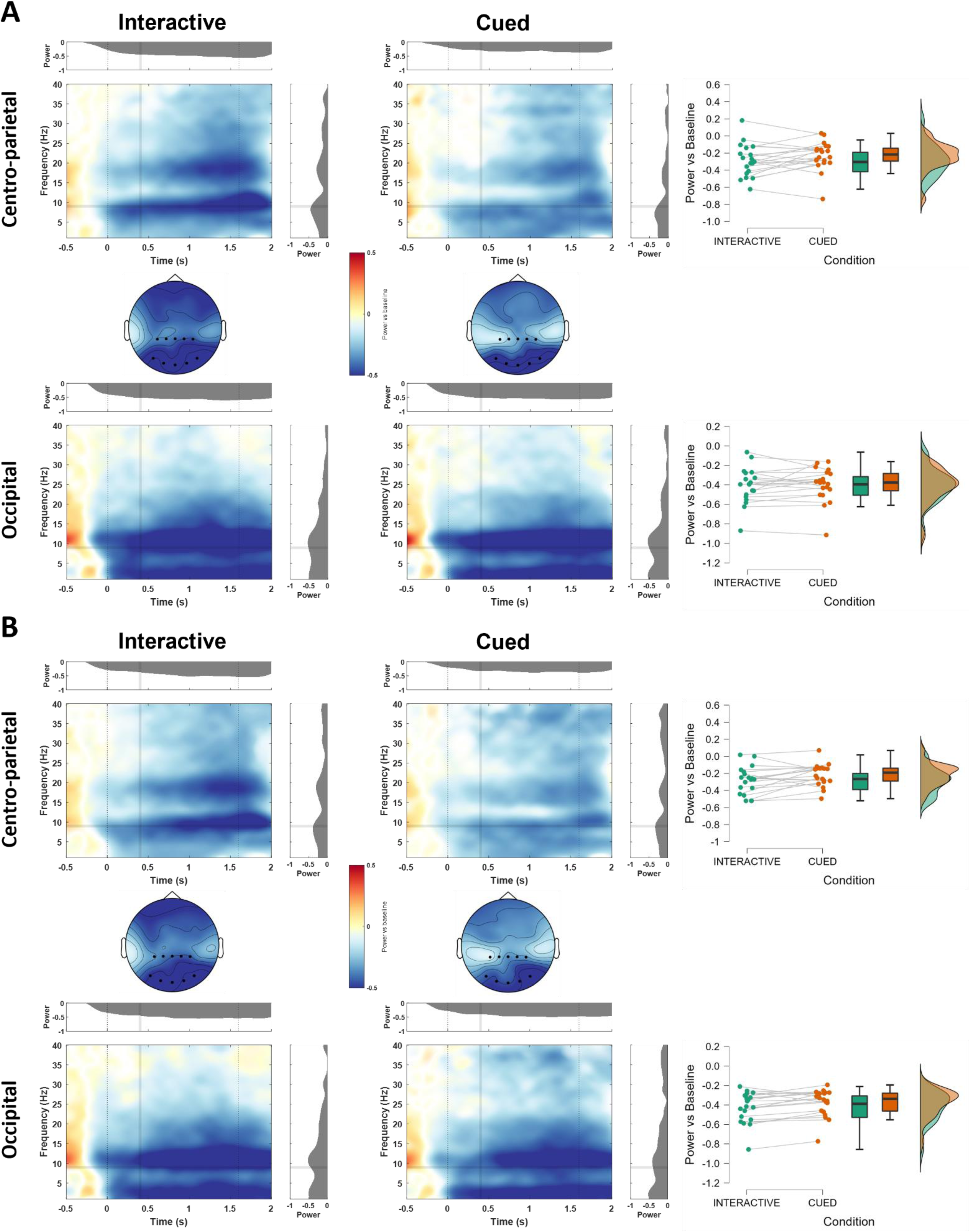
Power change relative to baseline over centro-parietal and occipital electrodes for all conditions (only NoCorrection trials). Panel A and B distinguish Body and NonBody trials, respectively. Topoplots in the middle and raincloud plots on the right show the mean Alpha (8-13 Hz) ERD in the time window between 0 (movement start) and 1600 ms (average movement end).

### Multivariate analysis

A Multivariate Pattern Analysis (MVPA) was performed to assess if and how a classifier would be able to distinguish the EEG patterns related to the observation and prediction of a movement performed by either a Body or a NonBody. The results are shown in Figures 6 and 7. In panel B of Figure 6 the results from the classification across time are reported, plotted in terms of percentage of accuracy, with statistically significant values of accuracy (*p* < .05, classification chance level 0.5) values of accuracy indicated by a bold line. The classifier was able to discriminate the appearance of the monitored interactor as early as ∼500 ms before movement start (significant clusters time window: [−600 −580 ms; −560 −410 ms]), with a first peak of classification accuracy around 250 ms before the stimuli (either body or dots) moved (i.e., time 0). Since all the videos began with the interactor being still for ∼1000 ms before starting to move, classification accuracy before 0 was likely driven by the visual processing of the interactor’s body shape. After the first peak, classification performance had a second peak around 100 ms after movement start and remained always significant until 700 ms after movement onset (significant cluster time window: [-390 700 ms]). The second peak was followed by a slow decay that remained significant for most of the movement period (significant clusters time window: [760 1090 ms; 1210 1550 ms; 1590 1640 ms]).

**Figure 6.**
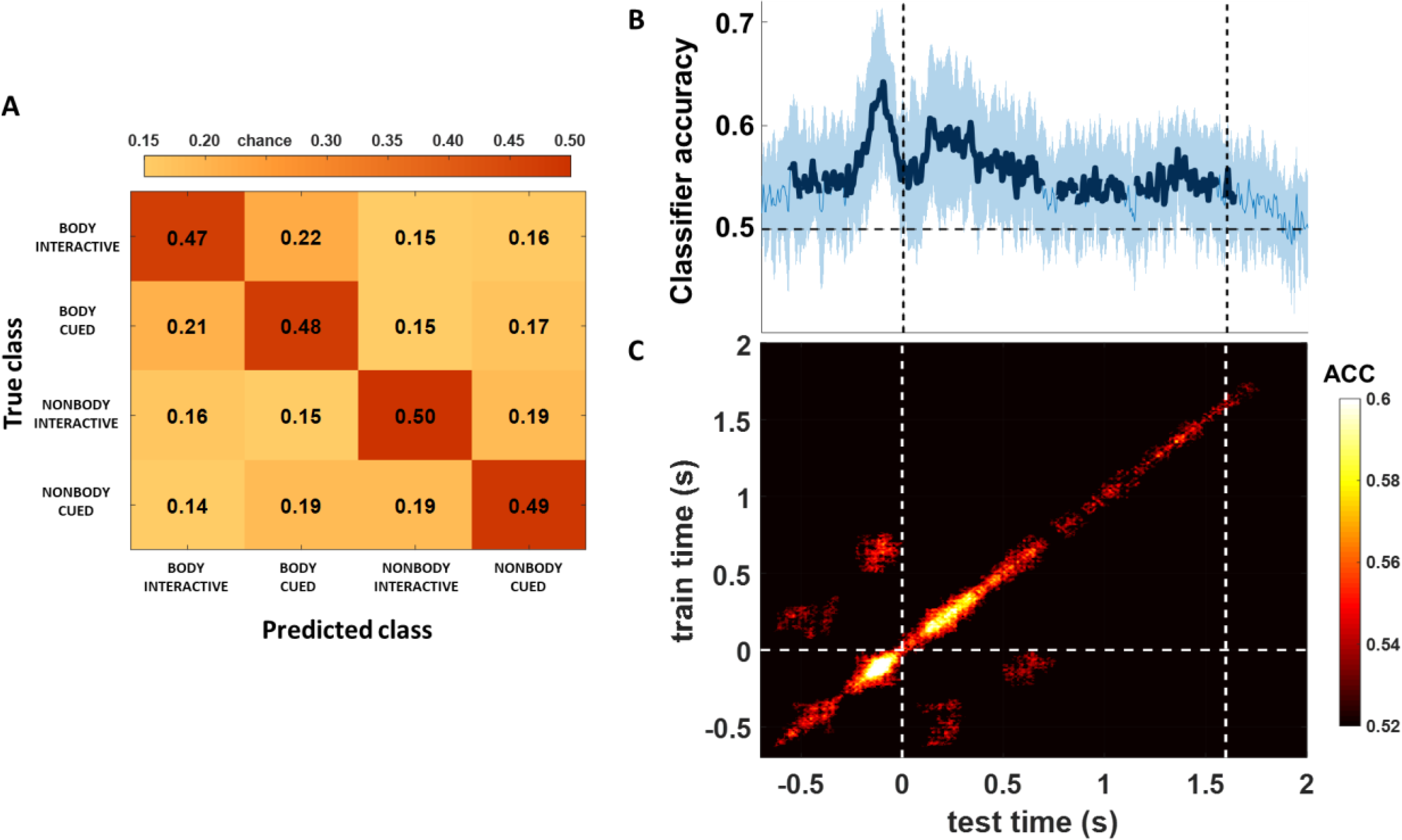
A) Confusion matrix showing the probability of the classifier (chance level 25%) to correctly predict the class of each sample in the movement time window (from 0 to 1600 ms). B) Classification scores across time, with significant (p < 0.05) accuracy clusters highlighted in bold. C) Temporal generalization matrix, with only significant accuracy scores (p < 0.05) plotted. Dashed lines on the x axis at 0 and 1.6 represent, respectively, movement start and average end time.

**Figure 7.**
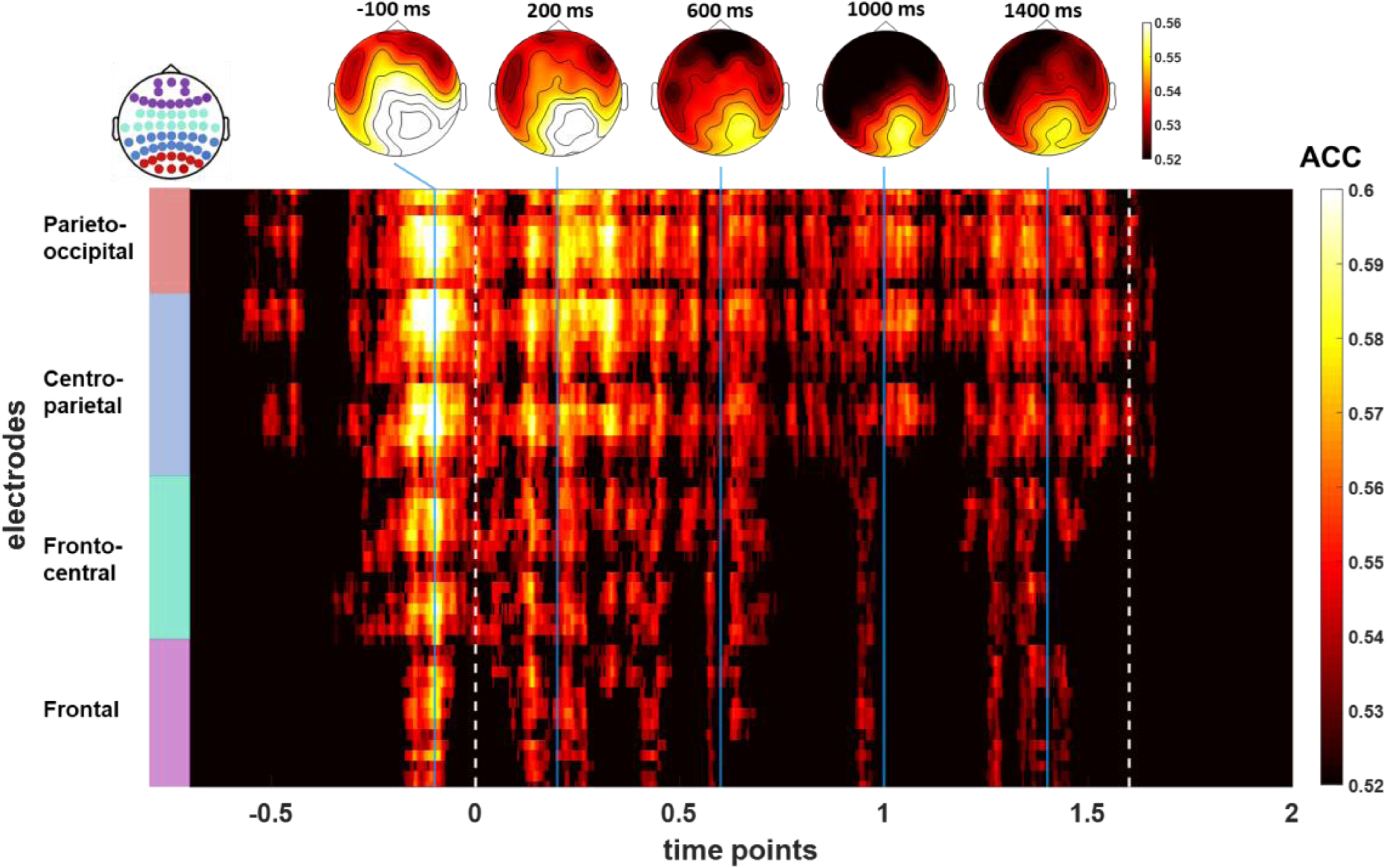
Searchlight analysis results plotted as a 2D matrix showing accuracy values for each electrode and time point (from −700 to 2000 ms around movement start), masked in order to show only points with significantly above chance level (p < .05) scores. The topographies above show accuracy values averaged in five time windows of interest. White dashed lines indicate the start of the movement (0 ms) and its average end (1600 ms).

Since we were interested in the neural patterns related to the observation of the movement (from 0 ms on), in order to investigate whether they differed from the patterns related to visual shape discrimination (occurring even before movement start) we ran a temporal generalization analysis. As proposed by (King & Dehaene, 2014), if a classifier generalizes from one time window to another, then the underlying processes for the two time windows would be similar, and the values in the resulting time x time plot would be represented as a single area spreading continuously out of the diagonal. If this is not the case, the results of the classification would be based on different information that travel across a chain of transient neural representations, and the results in the time x time matrix would yield a diagonal generalization pattern. The results of this approach are reported on Panel C of Figure 6. The time x time matrix, masked in order to show only significant results (*p* < .05), highlights a diagonal generalization pattern, suggesting that the neural processes related to the visual processing of a body form (likely occurring as early as the video started and before 0) are not the same as the ones related to the monitoring of a movement performed by a Body or a NonBody. Interestingly, however, we found two significant blobs at symmetrical points above and below the diagonal (around 200 and 600 ms). This suggests that a chain of transient representations (represented along the diagonal) where later generators involved in movement monitoring reactivate earlier ones (leading to symmetrical blobs outside the diagonal) involved in the processing of the interactor’s shape (King & Dehaene, 2014).

In order to test the weight of different electrodes to classification performance, we ran a searchlight analysis across time and electrodes with the same parameters and statistical analyses adopted for the classification across time points (i.e., LDA classifier, cluster-based permutation test correcting for multiple comparisons). The results are plotted in Figure 7 both on a 2D matrix of electrodes x time points and as topographies in five time-windows of interest (i.e., −100 ms, 200 ms, 600 ms, 1000 ms and 1400 ms from movement start). Overall, the searchlight decoding is in line with the results across time, highlighting a sustained contribution of right parieto-occipital electrodes to the classification from ∼300 ms before movement start until its average end (1.6 seconds).

Lastly, to investigate whether the neural patterns related to action observation were classifiable in terms of both appearance and the level of interactivity (Interactivity factor), we performed a multiclass classification across four experimental conditions (i.e., Body-Interactive, Body-Cued, NonBody-Interactive, NonBody-Cued). A multiclass Linear Discriminant Analysis (LDA) classifier was trained and tested on the EEG data in the time window between movement start (0 s) and movement end (1600 ms) for each condition. Figure 6A shows the resulting confusion matrix, allowing the assessment of the probability for the classifier to in/correctly discriminate between classes of data. Interestingly, the classifier showed a performance well above chance level (i.e., > .25) in distinguishing the four conditions (see the diagonal from top left to bottom right in the confusion matrix), as well as a higher tendency to confuse patterns related to same-appearance rather than same-condition factors (see lighter diagonal from bottom left to top right).

## DISCUSSION

In the present study we recorded EEG in participants engaged in a minimally interactive version of an ecological human-avatar joint-grasping task that has been previously adopted to study interactors’ action monitoring processes in different interactive contexts (Moreau et al., 2020). Specifically, we asked participants to observe a movement performed by either a Body or a (NonBody) control stimulus representing the same motion information (i.e., a set of dots with the very same human kinematics) and to synchronize their responses either only in time with the observed movement (i.e., Cued condition - subjects had to press a predefined key as synchronously as possible with the time when the body/dots touched to object) or in time and space (i.e., Interactive condition - subjects had to synchronize in time and to base their response depending on where the observed movement was directed, implying the necessity to continuously monitor the interactor’s movement).

Previous results from Gandolfo and colleagues (Gandolfo et al., 2019) tested whether visuo-motor interference effects were present when human participants were coordinating their actions either with a virtual avatar or with a stimulus conveying no visual information of the form of the body but complying with biological motion rules. Measuring the participants’ kinematic, they found no effect of the interactor’s body shape on visuo-motor interference. While these results did not highlight any specific activation for bodily vs non-bodily movements, a study where transcranial magnetic stimulation (TMS) was used to investigate the neurophysiological indices of motor excitability during motion observation (Agosta et al., 2016) found that, although the observation of both human movement and abstract motion modulated cortico-spinal excitability, only the former’s kinematics were significantly correlated with the activation of the observer’s motor cortex. Moreover, previous studies highlighted the role of the AMS in detecting discrepancies between sensory predictions and action outcomes during action observation and learning (Malfait et al., 2010) and during online interpersonal motor interactions (Boukarras et al., 2022; Moreau et al., 2020, 2022), as well as how this system is functionally related to posterior nodes of the AON at different levels depending on the interactivity of the context (Moreau et al., 2020). Here, we aim at further understanding whether the neural patterns associated with action observation (alpha/mu ERD/ERS) and action monitoring (i.e., oPe, oERN) are modulated by the bodily appearance of an interactor. Our results replicate previous ones on the activity of the action monitoring system and shed new light on the neural patterns related to action observation. Specifically, by combining univariate and multivariate analyses we show that: 1) among the electrocortical indices of action monitoring, only the early oPe was modulated by the bodily appearance of the interactor, whereas we found no trace of the oERN; 2) the classical desynchronization in alpha/mu (8-13 Hz) over central sites related to action observation was stronger when subjects had to predict in space and time the observed movement (i.e., Interactive condition), compared to when they only had to synchronize their choice in time (i.e., Cued condition); 3) posterior nodes of the AON continuously discriminate movements performed by a Body compared to a NonBody, and the neural patterns associated to such a process are also modulated by the degree of prediction required in order to fulfill the task requirements, even in a minimally interactive scenario.

### Interacting with a humanoid VP modulates the early Error Positivity (Pe)

Previous studies have shown how the time-dependent neural responses related to the activity of the action monitoring system are elicited both when people perform errors and observe another agent making errors (Somon et al., 2019; Spinelli et al., 2018; Zubarev & Parkkonen, 2018). Nevertheless, the oERN and the oPe have been associated with different processes occurring in the error-detection cascade. Specifically, the oERN is thought to index a fast comparison between an internally-generated prediction and incoming sensory inputs (Botvinick et al., 2001; Quilodran et al., 2008). Two sub-components of the oPe have been defined,anearly oPe and a late oPe, with the former occurring over frontocentral sites following the oERN and being associated predominantly to a reorientation response driven by the accumulation of evidence that an error has been committed/observed (Ridderinkhof et al., 2009; Steinhauser & Yeung, 2010), and the latter having a centroparietal topography and being related mainly to error awareness (Endrass et al., 2007; Ullsperger et al., 2014). Interestingly, Di Gregorio and colleagues (Di Gregorio et al., 2018) have studied how the Pe can occur in the absence of the ERN, thus pointing towards a functional dissociation between the systems underlying these two components.

In our study we report no trace of the oERN in any condition, whereas we report the occurrence of the early oPe only when a correction of the observed trajectory took place (Correction trials). Moreover, the amplitude of the early oPe was modulated by both the Appearance and the Interactivity factor, with larger early oPe in the Interactive condition compared to the Cued one as well as an increased amplitude for changes in the trajectory of a Body compared to a NonBody (i.e., only during Interactive trials).

With regard to the absence of the oERN in the present experiment, we hypothesize that this may be due to the minimal need for the subject to constantly anticipate and predict the observed movement, since participants observed the VP’s movement from a third-person perspective in order to eventually react to it. Thus, we hypothesize that the oERN is elicited when an early mismatch between an internally generated prediction and a change in the environment occurs, whereas in our paradigm subjects formed a minimal internal prediction leading to a complete absence of oERN.

Previous studies have shown how the amplitude of the early oPe is linearly modulated as a function of the observed error’s magnitude (Spinelli et al., 2018). Adding to this, our study highlights that such a top-down mechanism is modulated by the degree of prediction required during a minimally interactive interpersonal prediction task (i.e., larger amplitude for Interactive compared to Cued trials). Moreover, we report that, when a stronger attentional reorientation is needed (i.e., when subjects had to adapt their response based on the change in the movement’s trajectory inInteractive trials), the appearance of the interactor plays a role, with an increase in the early oPe amplitude for bodily movements compared to non-bodily ones. This result, together with our multivariate results, suggests that bodily movements are actually encoded differently than non-bodily ones at specific stages of the action observation and monitoring process.

With regard to the error-locked activity in the time-frequency domain, although previous results have shown that the rhythms in some nodes of the AON (i.e., LOTC) are in phase with the activation of the action monitoring system during social (human-avatar) interactions (Moreau et al., 2020, 2022), we found no evidence for a modulation of the midfrontal Theta rhythm by the appearance of the interacting partner (i.e., VP vs dots). Nevertheless, we replicated previous results highlighting a modulation of midfrontal Theta by top-down task-related processes (Interactivity factor).

### Centro-parietal alpha/mu desynchronization is modulated by top-down processes

The sensorimotor alpha/mu (8-13 Hz) desynchronization over central electrodes has often been the focus of EEG studies investigating action perception and understanding in interpersonal contexts (Fitzpatrick et al., 2019; Scanlon et al., 2022; Yun et al., 2012). Although our set-up was purely observational, lacking an action execution condition, our results broaden the literature on the role of alpha/mu suppression in the observation of actions in (minimally) interactive scenarios. Indeed, we report a significant difference between the alpha/mu desynchronization over central electrodes depending on the interactivity of the task. Specifically, we found a stronger decrease in power during the observation of a movement in the Interactive compared to the Cued condition. This result suggests a modulation of the activity in somatomotor areas by higher order, task-related processes, and are in line with previous studies showing a modulation of sensorimotor oscillations by the observation of different type of actions (Streltsova et al., 2010)), as well as studies where online interactions were investigated by means of dual-EEG (Ménoret et al., 2014). Notably, in line with previous studies (Urgen et al., 2013), we found no modulation of the alpha/mu suppression by the Appearance factor, suggesting that such rhythm, as well as midfrontal Theta, does not show any modulation for human appearances.

### The appearance of an interactor is coded continuously throughout its movement

Recent studies have shed light on the mechanisms underlying the perception of biological motion, showing how several cortical areas take part in this process at different hierarchical levels (Grosbras et al., 2012; Hirai & Senju, 2020; Puce & Perrett, 2003; Saygin et al., 2012; Troje & Chang, 2023; Van Overwalle, 2009). For example, Duarte and colleagues (Duarte et al., 2022) reported that during the observation of point lights representing the main joints of a person and moving in accordance to biological motion kinematics rules, at least two different stages of motion processing must be differentiated. Specifically, the brain must distinguish motion patterns at the level of each individual dot (i.e., local motion) and integrate them into the percept of a coherent body where these multiple joints bear together (i.e., global motion). The authors investigated how these two patterns are serially processed in specific areas of the LOTC by means of functional magnetic resonance imaging (fMRI) and multivariate analyses. Their results highlighted a two-stage framework for the neural processing of biological motion, with early (i.e., hMT+ and V3A) and higher-level (i.e., Fusiform Gyrus - FFG - and EBA) visual areas processing local motion patterns, whereas the posterior STS (pSTS) was involved in global motion processing. Their results provide strong, highly spatially resolved evidence that pairs with another recent study from Chang and colleagues (Chang et al., 2021), where the authors characterized the spatiotemporal characteristics of the perception of biological motion by means of magnetoencephalography (MEG). In detail, through a combination of univariate and multivariate analyses, they showed that the neural patterns associated with global motion processing can be discriminated from those related to local kinematics as early as 100 ms after the beginning of the point-light display movement, and that such discrimination remains relatively stable throughout the majority of the movement period.

Our results broaden the above-mentioned ones, by investigating whether the neural activity generated by the predictive observation of bodily movements performed by a VP (instead of point lights) may be distinguished by the neural activity associated to the observation of the movements of a set of dots and lines showing the very same kinematics. Our stimuli allowed to further understand, the influence of top-down processes over these activities, as participants were asked to observe the movements while being asked to predict their spatio-temporal development at different degrees in cued and interactive conditions. We report that a LDA classifier applied on participants’ EEG signal can successfully decode bodily from non-bodily movements as early as ∼100 ms after movement start (in line with Chang and colleagues’ results), and that such significant decoding, after a slight decay, remains relatively stable throughout the entire movement period. Moreover, by performing a temporal generalization analysis, we were able to show that the neural patterns contributing to such classification are not the same related to the processing of a still body shape, although the generators contributing to such discrimination are recalled in early time windows during bodily-movement decoding. Furthermore, our searchlight classification analysis revealed the electrodes the classifier relied the most on to decode the different stimuli, highlighting a sustained contribution of mainly right occipito-parietal electrodes throughout the entire movement period, suggesting that the neural patterns from these areas are crucial for the discrimination of global bodily movements. Our results thus broaden the understanding of the role of the occipito-parietal node of the AON, following previous studies that highlighted how it is crucial for discriminating between the meaning and the effector of an action (Lingnau & Petris, 2013), recognizing different actions (Urgesi et al., 2014), while also having an early access to abstract action representations (Moreau et al., 2023; Tucciarelli et al., 2015). Our results are also in line with previous fMRI ones reporting right lateral occipito-temporal areas as a crucial node for global motion processing (Duarte et al., 2022; Sokolov et al., 2018).

Lastly, the results from our multiclass classification analysis strengthened the ones from the binary (i.e., Body vs NonBody) classification, showing that our classifier was able to distinguish the appearance of the observed agent while also classifying correctly trials in the Interactive condition from trials in the Cued one. Interestingly, the classifier tended to confuse more same-appearance samples rather than same-condition ones, suggesting that the appearance of the observed agent was clearly distinguishable even across different tasks.

## CONCLUSIONS

By means of EEG, we investigated if and how biological movements are processed differently based on their bodily/non-bodily appearance in a minimally interactive scenario. We targeted the neural correlates of the activity in different nodes of the AON as well as in the action monitoring system. We found that the sensorimotor alpha/mu rhythm typically related to action observation is modulated by top-down, task-related processes, with a stronger desynchronization when subjects had to rely on spatial and temporal features of the observed movement (i.e., predict and monitor the target of the observed movements) compared to when they only had to synchronize their response to the stimuli, whereas we found no modulation by the appearance of the agent moving. Rather, our multivariate results showed that such a feature is related to specific, parieto-occipital neural patterns that remain stable throughout the entire duration of an observed movement. We showed that such patterns differ from the ones related to initial shape processing (i.e., differentiating a still body shape from a non-body one), although the two overlap at specific stages during movement observation, suggesting a reactivation of the generators related to body-shape processing at later times. Lastly, we replicated previous results showing a modulation of the electrocortical indices of error monitoring by top-down, task related processes, while only the early oPe was responsive also to the appearance of the observed agent, depending on the task. Taken together, these findings broaden our understanding of the influence of bodily appearance on the spatiotemporal processing of biological movements in the AON and in the action monitoring system during (minimally) interactive scenarios. These results may potentially inform advancements in the field of human-artificial-agent interaction, fostering the development of technologies that align with the complexities of neural processing during social interactions.

